# Unbiased, systematic surfaceome profiling of Acute Leukemia to identify novel immunotherapy targets

**DOI:** 10.64898/2026.09.16.752145

**Authors:** Benson M. George, Anna Rodriguez-Pöhnlein, Donavelle J. DeCastro, Jennifer Porat, Benyu Zhou, Dinah M. Abeja, Ryan J. Lumpkin, Rebecca J. Metivier, Anna Schmoker, Wencke Walter, Torsten Haferlach, Yana Pikman, Katherine A. Donovan, Kimberly Stegmaier, Eric S. Fischer, Franziska Wachter, Ryan A. Flynn

## Abstract

Cell surface proteomics provides a direct topological assessment of the outer membrane of cells and enables the capture of low abundance proteins that may be missed by whole cell proteomics. Here we present an unbiased atlas of the whole cell and surface proteomes of 25 commonly used leukemic cell lines, encompassing both lymphoid and myeloid lineages, and a variety of driver mutations. Paired-wise analysis highlights recurrent surface proteins that are not detected by whole cell proteomics. Coupling this dataset to RNA-sequencing, we also discovered genes where protein and RNA abundances are discordant. In KMT2A-rearranged AML, CD70 expression was increased across cell lines and validated in primary patient samples, supporting CD70 as a candidate therapeutic target in this disease. Several proteins are enriched in the surface proteomes but lack surface annotation, adding to the list of non-canonical cell surface proteins, including RNA-binding proteins, an emerging class of candidate immunotherapeutic targets.

**Statement of Significance:** This mass spectrometry atlas pairs whole-cell and cell-surface proteomes across 25 leukemia cell lines, creating a resource for understanding these widely used models. Direct surface profiling captures proteins missed by transcriptomic and whole-proteome data, enabling interrogation of cell interactions and nomination of immunotherapy targets, exemplified by CD70 in KMT2A-rearranged AML.

## Introduction

Acute myeloid leukemia (AML) remains a high-risk malignancy, with overall survival of approximately 35% in adults and 65% in children. Outcomes are particularly poor for patients with relapsed or refractory disease. Although graft-versus-leukemia effects after allogeneic hematopoietic stem cell transplantation demonstrate that AML can be immunologically controlled, effective immunotherapeutic strategies remain limited, with gemtuzumab ozogamicin representing the only established surface antigen–directed approach^1–3^. A major limitation in developing surface protein targeted therapies in AML has been the lack of identified antigens that are differentially expressed on malignant blasts, and absent on vital tissues, like hematopoietic stem cells.

This situation is countered by the experience in acute lymphoblastic leukemia (ALL), where lineage associated proteins like CD19, CD20 and CD22, have enabled the use of monoclonal antibodies, antibody-drug conjugates (ADC), bispecific T-cell engagers (TCE), and chimeric antigen receptor T-cells (CAR-Ts)^4–7^. Though these markers are expressed on healthy B-cells, given their relatively expendable nature, therapies targeting these have enabled significant advances in the treatment of B-cell malignancies^6,7^. However, as these therapies become more widely used and are integrated into treatment regimens for patients with newly diagnosed leukemias, there is a critical need to identify additional targetable antigens for immunotherapies that can be used in patients with relapsed or refractory leukemias.

Cell surface proteins are attractive therapeutic targets in leukemia because they can be directly engaged by ADCs, TCE, or CAR-Ts. However, target nomination has often relied on flow cytometry, transcriptomic profiling, or whole-proteome datasets, which all have limitations. Flow cytometry is limited to interrogated only predefined antigens. Transcriptomic profiling does not reliably predict total or surface protein levels^8^. Whole-proteome analysis may miss low-abundance or selectively surface-enriched proteins. Comprehensive proteome analysis of the healthy bone marrow has been performed previously but without specific surface protein (surfaceome) enrichment.^9^ These limitations create a critical gap in leukemia target discovery, particularly in pediatric AML, where lineage- and subtype-specific surface vulnerabilities remain incompletely defined.

Prior AML surfaceome studies^10–15^ have established the utility of cell surface proteomics in primary AML samples, including large adult AML cohorts. However, limited material, variable blast purity, prior treatment exposure, and patient-to-patient heterogeneity, can limit the utility of this work. Profiling cell lines enables controlled and reproducible comparison of antigen expression across genetically defined subgroups, which is difficult to achieve in primary samples. Further, coupling this data with RNA-sequencing and whole cell proteomics can enable broader biological understanding of dynamics in translation and surface localization.

To address this gap, we performed unbiased cell surface proteomic profiling of 25 leukemia cell lines with matched whole-proteome and RNA sequencing data. This approach identified CD70 as significantly enriched in *KMT2A*-rearranged AML cell lines, a finding we subsequently validated in primary patient samples. Additionally, surface enrichment provided a more comprehensive view of the surface proteome, capturing numerous non-canonical surface proteins, such as RNA binding proteins.

## Results

### Paired whole cell and surface mass spectrometry of leukemic cell lines

We performed whole cell and surface enrichment mass spectrometry of 25 commonly used human leukemia cell lines (**Table 1**) spanning lymphoid and myeloid lineages and a wide spectrum of driver mutations. The most common genetic abnormalities across all cell lines were *KMT2A* rearrangements, found in 7/20 myeloid cell lines and 1/5 lymphoid cell lines, and *TP53,* seen in 13 cell lines.

**Table 1.**
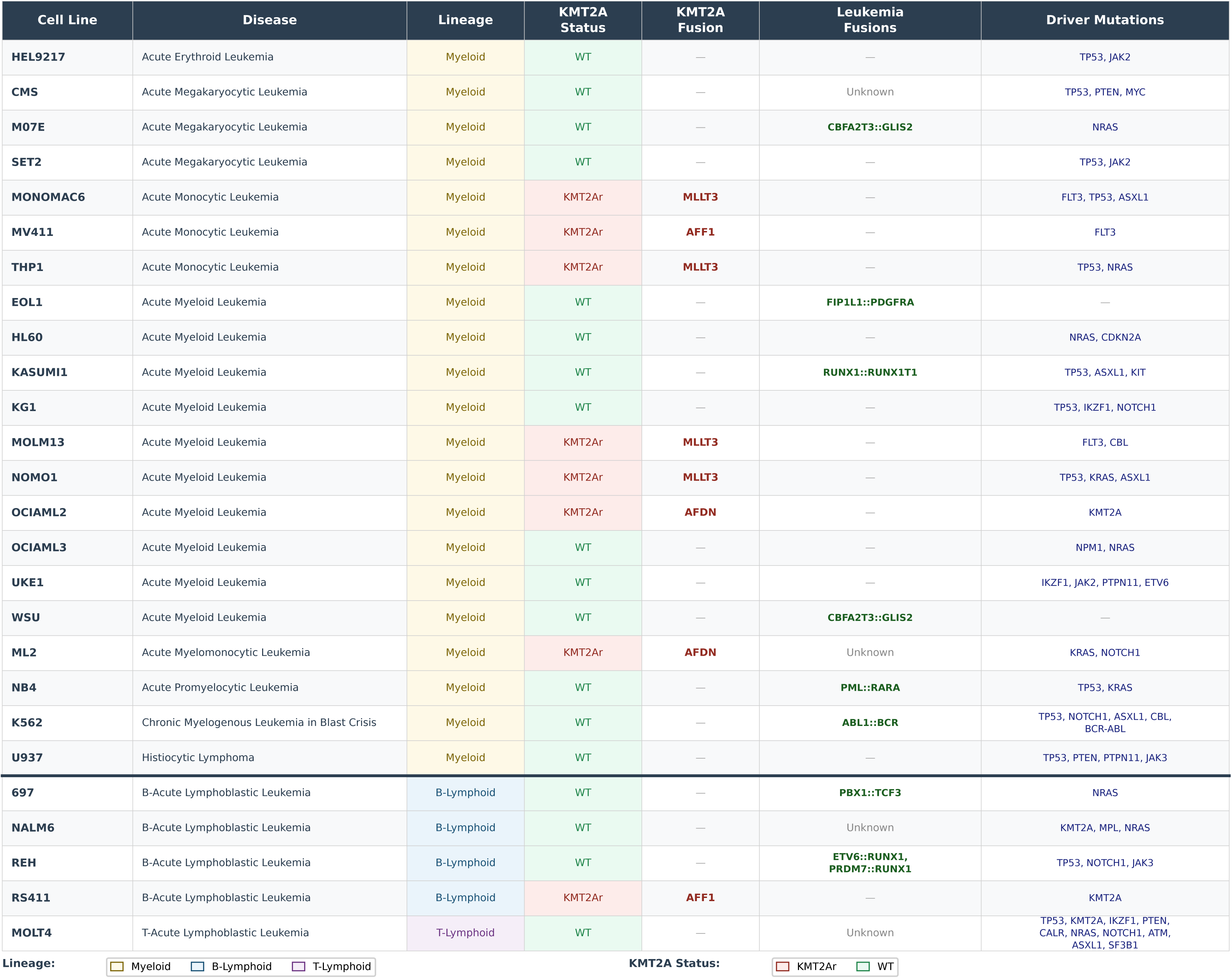
Cell lines analyzed and annotated to reflect hematopoietic lineage, KMT2A rearrangement status, KMT2A fusion partner, other reported fusions and driver mutations.

| Cell Line | Disease | Lineage | KMT2A Status | KMT2A Fusion | Leukemia Fusions | Driver Mutations |
| --- | --- | --- | --- | --- | --- | --- |
| HEL9217 | Acute Erythroid Leukemia | Myeloid | WT | — | — | TP53, JAK2 |
| CMS | Acute Megakaryocytic Leukemia | Myeloid | WT | — | Unknown | TP53, PTEN, MYC |
| M07E | Acute Megakaryocytic Leukemia | Myeloid | WT | — | CBFA2T3::GLIS2 | NRAS |
| SET2 | Acute Megakaryocytic Leukemia | Myeloid | WT | — | — | TP53, JAK2 |
| MONOMAC6 | Acute Monocytic Leukemia | Myeloid | KMT2Ar | MLLT3 | — | FLT3, TP53, ASXL1 |
| MV411 | Acute Monocytic Leukemia | Myeloid | KMT2Ar | AFF1 | — | FLT3 |
| THP1 | Acute Monocytic Leukemia | Myeloid | KMT2Ar | MLLT3 | — | TP53, NRAS |
| EOL1 | Acute Myeloid Leukemia | Myeloid | WT | — | FIP1L1::PDGFRA | — |
| HL60 | Acute Myeloid Leukemia | Myeloid | WT | — | — | NRAS, CDKN2A |
| KASUMI1 | Acute Myeloid Leukemia | Myeloid | WT | — | RUNX1::RUNX1T1 | TP53, ASXL1, KIT |
| KG1 | Acute Myeloid Leukemia | Myeloid | WT | — | — | TP53, IKZF1, NOTCH1 |
| MOLM13 | Acute Myeloid Leukemia | Myeloid | KMT2Ar | MLLT3 | — | FLT3, CBL |
| NOMO1 | Acute Myeloid Leukemia | Myeloid | KMT2Ar | MLLT3 | — | TP53, KRAS, ASXL1 |
| OCIAML2 | Acute Myeloid Leukemia | Myeloid | KMT2Ar | AFDN | — | KMT2A |
| OCIAML3 | Acute Myeloid Leukemia | Myeloid | WT | — | — | NPM1, NRAS |
| UKE1 | Acute Myeloid Leukemia | Myeloid | WT | — | — | IKZF1, JAK2, PTPN11, ETV6 |
| WSU | Acute Myeloid Leukemia | Myeloid | WT | — | CBFA2T3::GLIS2 | — |
| ML2 | Acute Myelomonocytic Leukemia | Myeloid | KMT2Ar | AFDN | Unknown | KRAS, NOTCH1 |
| NB4 | Acute Promyelocytic Leukemia | Myeloid | WT | — | PML::RARA | TP53, KRAS |
| K562 | Chronic Myelogenous Leukemia in Blast Crisis | Myeloid | WT | — | ABL1::BCR | TP53, NOTCH1, ASXL1, CBL, BCR-ABL |
| U937 | Histiocytic Lymphoma | Myeloid | WT | — | — | TP53, PTEN, PTPN11, JAK3 |
| 697 | B-Acute Lymphoblastic Leukemia | B-Lymphoid | WT | — | PBX1::TCF3 | NRAS |
| NALM6 | B-Acute Lymphoblastic Leukemia | B-Lymphoid | WT | — | Unknown | KMT2A, MPL, NRAS |
| REH | B-Acute Lymphoblastic Leukemia | B-Lymphoid | WT | — | ETV6::RUNX1, PRDM7::RUNX1 | TP53, NOTCH1, JAK3 |
| RS411 | B-Acute Lymphoblastic Leukemia | B-Lymphoid | KMT2Ar | AFF1 | — | KMT2A |
| MOLT4 | T-Acute Lymphoblastic Leukemia | T-Lymphoid | WT | — | Unknown | TP53, KMT2A, IKZF1, PTEN, CALR, NRAS, NOTCH1, ATM, ASXL1, SF3B1 |

Lineage:
Myeloid
B-Lymphoid
T-Lymphoid
KMT2A Status:
KMT2Ar

Amongst all cell lines, the average number of unique proteins detected by whole cell proteomics was 5,009 per line, while surface proteomics defined an average of 682 unique proteins. To perform an initial classification of the surface-enriched proteins, we intersected our hits with an *in silico* surface prediction score (SPC)^16^, where the score is the sum of predictive databases in which a particular protein is postulated to be on the cell surface. For example, a score of 4 corresponds to a higher likelihood of being on the cell surface, as it indicates that a given protein is found in all four separate databases. Proteins with no surface prediction (SPC=0) made up 86.4% of the whole cell proteome on average and 55.8% of the surface proteome, while SPC=4 proteins represented 1.5% of the whole cell proteome and 39.8% of surface proteome (**Figure 1A, Supplemental Figure 1**). The increased representation of SPC=4 proteins detected in surfaceomes as compared to whole cell proteomes validates our enrichment strategy.

**Figure 1:**
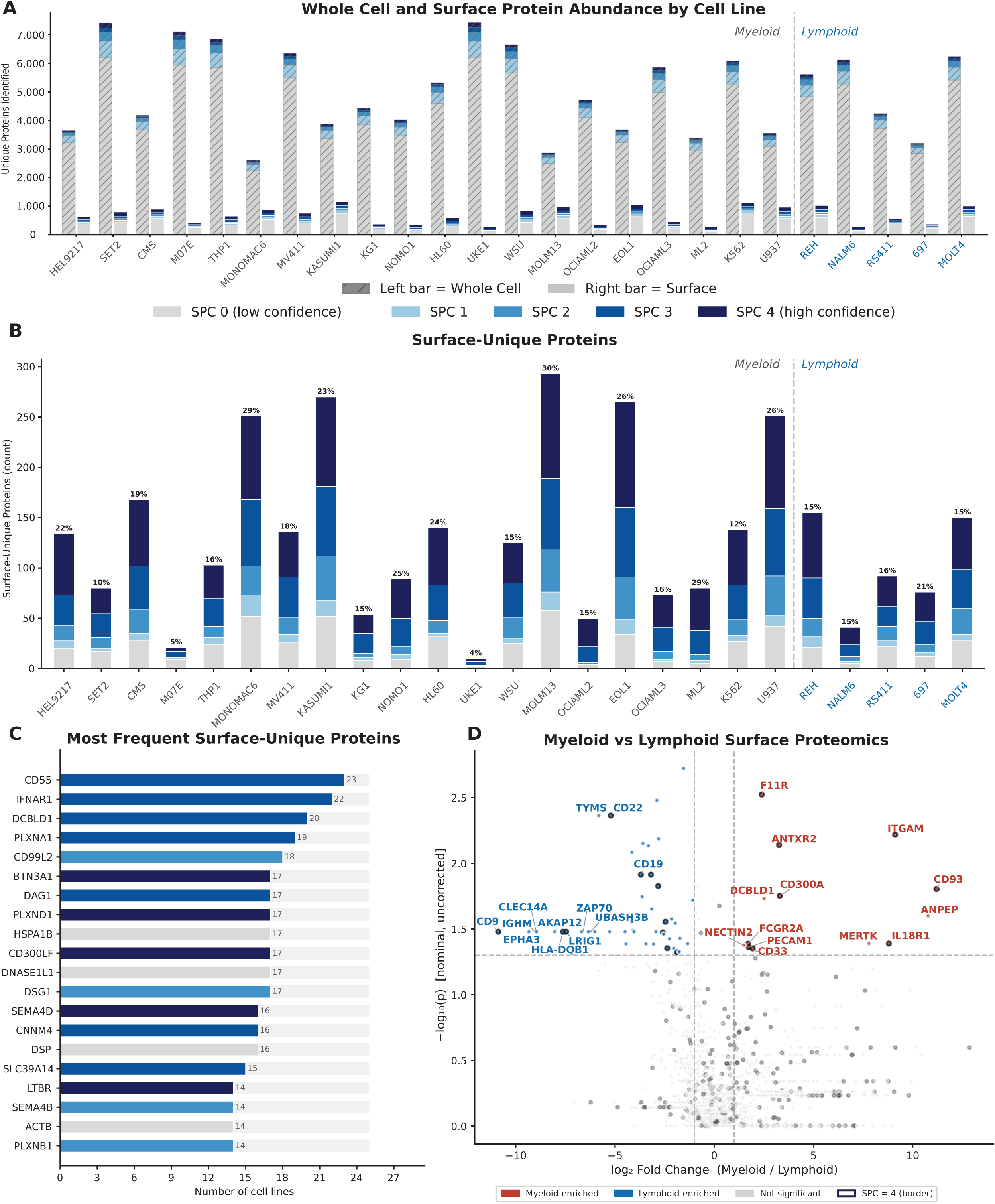
An atlas of proteomes across 25 leukemic cell lines. A. Total number of unique proteins detected in whole cell and surface proteomics for each cell line. Surface proteome is denoted by stacked bars showing surface prediction score (SPC). B. Stacked bar plot of total number of proteins found in surface proteomics but not found in whole cell proteomics for each cell line with colors corresponding to SPC annotation. C. Proteins most frequently found only in surface proteomics across all cell lines with colors corresponding to SPC annotation. D. Volcano plot showing differentially abundant proteins discovered in surface proteomics dataset comparing myeloid (red) and lymphoid (blue) cell lines.

Whole cell proteomics can fail to identify known cell surface proteins if their overall abundance in the cell is low. Our dataset highlights the utility of surface enrichment by the identification of proteins that are exclusively detected in the surface proteome (**Figure 1B**). On average, we see 135 surface-enriched proteins per cell line that are never detected in paired whole cell proteomics; this is ∼25%, on average, of the empirically captured surface proteome. When examining proteins only discovered via surface-enrichment, 25 of these are identified in more than half (13) of the cell lines assayed (**Figure 1C**). CD55, a glycophosphatidylinositol (GPI) anchor protein involved in the complement pathway was found on 23 of 25 cell lines and consistently present only in surface enriched datasets across both myeloid (**Figure S2A**) and lymphoid (**Figure S2B**) cell lines.

Conversely, we looked for proteins predicted to be on the surface, which were present in whole cell datasets, but not surface enriched proteomes. We uncovered that a small fraction of each whole-cell proteome, typically <1%, consisted of SPC4 proteins that were not detected in the corresponding surface-enriched proteome (**Figure S2C**), including CD46, CD151, and HLA-E. Some of these proteins are recurrently absent from surface proteomes (**Figure S2D**), so we examined if there could be a biochemical rationale for this finding. We used sulfo-NHS-SS-Biotin as our cell surface biotinylation reagent, which relies upon the availability of surface lysines on the target protein for conjugation. Therefore, we examined lysine abundance among SPC=4 proteins that were not detected in the corresponding surface-enriched proteome. We found that the average number of lysines (**Figure S3A**), length of the extracellular domain (ECD) (**Figure S3B**), and number of lysines within the ECD (**Figure S3C**) for these proteins was significantly lower than surface predicted proteins that were detected in surfaceomes. The density of lysines within the ECD (**FigureS3D**) did not seem to impact enrichment. These data highlight that while surface-enrichment facilitates the discovery of surface-expressed proteins that are difficult to detect in bulk proteomics data, these methods may be biased by the enrichment chemistry used.

To further validate our dataset, we compared myeloid and lymphoid cell lines and observed the expected lineage-defining expression patterns. For example, we see that PAX5, which is a B-cell transcription factor, is upregulated in B-lymphoid lines. Similarly, we see ANPEP (CD13), a known myeloid marker, associated with myeloid cell lines (**Figure S2E**). The surface proteomes further separated these cohorts with other canonical lineage receptors (**Figure 1D**), such as CD22 and CD19 for B-lymphoid lines.

Several of the differentially expressed cell-surface proteins identified in our analysis, including EPHA3 and CD9, are currently being investigated as novel therapeutic targets in B-ALL^17–19^. We additionally found receptors that have not been previously reported in these subsets, such as LRIG1 (**Figure 1D**). This receptor was found only on 697 and NALM6 cell lines, however, LRIG1 has mostly been described on epidermal stem cells and T-regulatory cells^20,21^.

This dataset also offered a unique opportunity to examine WSU-AML and M07e, which are both originally derived from the same patient sample^22^. Though these cell lines have drifted by mutational analysis (**Supplemental Figure 4A**), they are far more similar to one another by proteomics than any other pair of cell lines (**Supplemental Figure 4B**), including the four myeloid cell lines sharing the same *KMT2A-MLLT3* driver fusion. Despite this, there are still numerous differentially expressed proteins (**Supplemental Figure 4C-D**), such as the entire MHC Class 1 complex (B2M, HLA-A, HLA-B, HLA-C), which is nearly absent from WSU-AML (**Supplemental Figure 4E-F**). As there are no reported mutations in B2M or HLA Class I genes in WSU-AML, this cell line could be used as a useful model to study non-canonical downregulation of MHC.

### Correlation between transcriptomes and proteomes

Twenty-two cell lines analyzed in this dataset have corresponding RNA sequencing data available through the Cancer Cell Line Encyclopedia (CCLE)^23^, providing an opportunity to correlate steady state transcriptomes and proteomes. Each cell line showed a direct relationship between transcript and protein abundance (**Supplemental Figure 5**). The dynamic range was narrower for proteomics than RNA-sequencing, as previously reported and expected^24^. Across all cell lines, the relationship between protein and RNA abundance was quite consistent with a Spearman correlation coefficient range from 0.46 to 0.59 (**Supplemental Figure 5**).

When aggregating the data from all 22 cell lines, we created Z-scores for both RNA and protein to understand their relationship at scale. The Z-scores provide a standardized measure of each gene’s relative RNA and protein abundance. Across genes, RNA and protein abundance showed an overall positive relationship. Importantly, this enabled identification of outliers in which RNA and protein abundance for a given gene were disproportionately discordant (**Figure 2A**). For example, we identified multiple histone proteins routinely ranked higher by protein abundance than RNA abundance. Conversely, we found that mitochondrial proteins often had a far higher RNA abundance rank than protein rank. These outlier relationships could serve as useful experimental models to explore protein degradation and RNA half-life dynamics.

**Figure 2:**
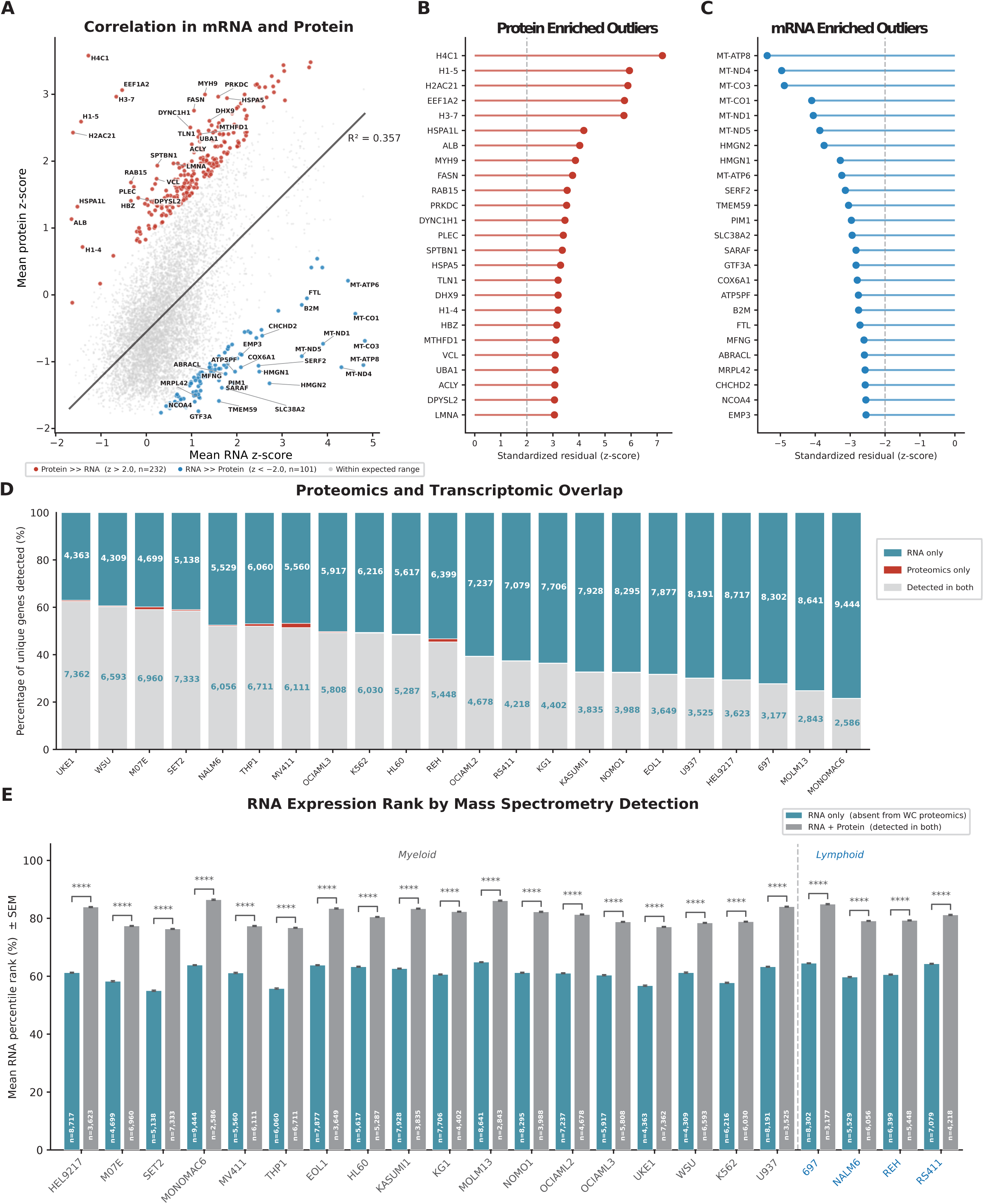
Unified comparison of transcriptomes and proteomes. A. Scatter plot of mean protein abundance (log10 PPM+1) vs mean mRNA expression (log2 TPM+1, CCLE) across 7,660 genes detected in ≥3 matched cell lines, with OLS regression (R²=0.349). Genes with |z| > 2.0 are classified as discordant, with 173 protein-enriched (z > 2.0, blue) and 135 mRNA-enriched (z < −2.0, red) outliers. B-C. Flanking lollipop charts show the top 25 outliers in each direction by standardized residual z-score. D. Stacked bar chart showing per-cell-line gene detection overlap between whole-cell proteomics (mean PPM > 0) and CCLE mRNA (log2 TPM+1 > 0.5); bar annotations show absolute gene counts for each category. E. Mean log2(TPM+1) ± SEM of RNA-only genes (RNA log2 TPM+1 > 0.5; proteomics mean PPM=0) vs RNA+Protein genes (detected in both, RNA log2 TPM+1 > 0.5; proteomics mean PPM>0) per cell line, **** FDR-adjusted Mann-Whitney U p < 0.0001.

Next, we characterized the relative detection sensitivity of the proteomics and RNA sequencing datasets. Across all cell lines, we discovered that an average of 4955 genes were detected in both the transcriptome and proteome (**Figure 2D**). There was an average of 108 genes found by proteomics only and 6838 genes identified by RNA-sequencing only. To further characterize the greater detection sensitivity of RNA sequencing, we compared transcript abundance between genes detected exclusively by RNA sequencing and those detected by both RNA sequencing and mass spectrometry. Genes detected in both datasets had relatively high transcript abundance, with mean transcripts per million (TPM) values ranking between the 77th and 87th percentiles, whereas genes detected only by RNA sequencing ranked between the 55th and 65th percentiles **(Figure 2E, supplemental Figure 6A).** Using this data, we modeled the probability of detection by whole cell proteomics based on RNA sequencing output and found that transcripts in the 75^th^ percentile resulted in a 50% probability of that protein being detected by mass spectrometry. On a per-cell-line basis, this ranged between 60^th^ and 90^th^ percentile (**Supplemental Figure 6C**), which could be related to cell-dependent translation efficiency or RNA turnover. These metrics provide a quantitative boundary to leverage RNA sequencing data as it would relate to the translated proteome.

### Proteomic Profile of *KMT2A*r Leukemia

*KMT2A* rearrangements (*KMT2Ar)* comprise approximately 3–6% of adult and 15–20% of pediatric AML and define a clinically high-risk group characterized by substantial relapse risk, although prognosis varies according to the specific KMT2A fusion partner ^25–27^. With multiple cell lines driven by a *KMT2Ar*, we sought to determine whether there was a characteristic proteomic signature as compared to *KMT2A*-wild type (*KMT2*Awt) AML lines. 72 proteins in the whole cell proteome and 30 in the surface enriched proteome were predominantly associated with *KMT2Ar* myeloid cell lines (**Figure 3A-B**). We compared this with the CCLE transcriptome data for these lines and discovered that only 35.9% of *KMT2Ar*-enriched genes by proteomics were also significantly different by transcriptomics (**Figure 3C**). This again highlights how RNA sequencing alone can fail to serve as a surrogate for protein expression in some instances. We next correlated the proteomics findings with primary patient data. Using the BEAT-AML cohort^28^, we tested whether *KMT2Ar* surfaceomics hits were differentially expressed in patient sample transcriptomes. Of the 30 top surfaceomics hits, 29 had transcriptome data, and 21 of 29 were enriched in the BEAT-AML cohort (**Figure 3D**). Taken together, proteomic analysis nominates *KMT2A*r-specfic pathways that also appear to be subtype specific in *KMT2A*r primary patient samples.

**Figure 3:**
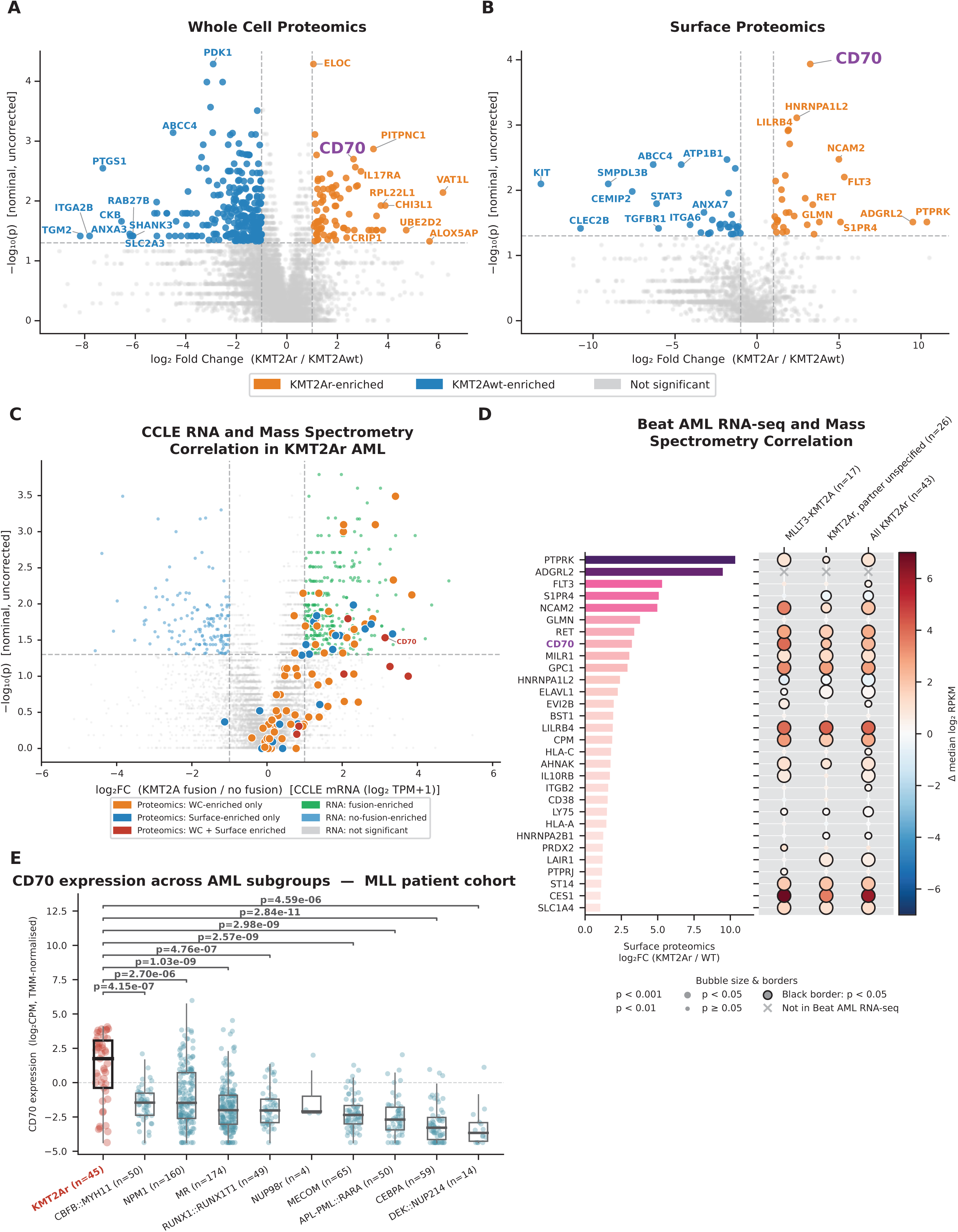
Integrated expression analysis nominates CD70 as an upregulated protein in KMT2Ar AML. A-B. Volcano plots comparing KMT2A-rearranged (KMT2Ar) vs KMT2A wild-type (KMT2Awt) myeloid cell lines in whole-cell (A) and surface (B) proteomics, using two-sided Mann-Whitney U with a 1 PPM pseudocount for log2 fold change. Significant hits (nominal p < 0.05, |log2FC| > 1.0, uncorrected) are highlighted, with the top 10 enriched proteins by Manhattan distance are labeled for each group of cell lines. C. Volcano plot of CCLE mRNA expression (log2 TPM+1) comparing KMT2A-fusion (n=6) vs no-fusion (n=11) AML cell lines restricted to proteomics-matched lines, assessed by two-sided Mann-Whitney U (nominal p < 0.05, |log2FC| > 1.0). KMT2Ar-enriched proteomics hits from the whole-cell and/or surface analysis are overlaid as larger colored points. D. Most enriched surface proteins in KMT2A-r AML cell lines ranked in order of fold enrichment over non-rearranged cell lines. Bubble color shows Δ median log2 RPKM (red = KMT2Ar-enriched, blue = depleted) amongst BEAT-AML KMT2Ar subgroups, bubble size reflects p-value, black borders indicate p < 0.05, and × marks genes absent from the Beat AML RNA-seq dataset. E. Comparison of CD70 expression obtained from Munich Leukemia Laboratory (MLL) patient samples across various AML subpopulations and compared to KMT2Ar samples (red). P-values above brackets are from two-sided Mann-Whitney U tests comparing each subgroup against KMT2A (FDR-BH corrected).

Of *KMT2A*r-associated proteins, CD70 emerged as a candidate target of particular interest. Although physiologic CD70 expression is largely restricted to activated immune cells, aberrant expression has been demonstrated on AML blasts and leukemia stem cells. CD70-directed therapeutics have consequently been developed across multiple modalities, including monoclonal antibodies, ADCs, and CAR-T cells, with anti-CD70 antibodies and cellular therapies advancing to clinical evaluation in AML^29–32^. In a phase 2 trial combining anti-CD70 with azacitadine^33^, there were modest responses. However, in light of the molecular heterogeneity of AML, it is possible that selecting for particular genetic subsets could yield more robust responses. Therefore, we next focused on CD70, which was enriched in *KMT2Ar* myeloid cell lines by proteomics and transcriptomics. Using RNA sequencing data from 670 AML samples acquired by the Munich Leukemia Laboratory, we tested CD70 expression across various genetic subtypes. CD70 expression was statistically significantly upregulated in *KMT2Ar* patients when compared to every other genotype (**Figure 3E**). *KMT2A* has many known fusion partners; *KMT2A*r myeloid cell lines included in our dataset were fused to *MLLT3* (n=3), *AFDN* (n=2), and *AFF1* (n=1). Among primary patient samples, MLLT3, MLLT10, and MLLT1 fusions were associated with significantly higher CD70 expression than KMT2A-wild-type AML (**Supplemental Figure 7**). *ELL*, *MLLT11*, and *AFDN* samples had similar expression to *KMT2A*wt AML, suggesting that fusion partner may influence CD70 expression. Taken together, this data highlights the need to do a focused clinical trial of CD70 targeting therapies in *KMT2A*-rearranged AML, where there are likely to be more patients with higher expression.

### Non-Canonical Surface Proteins

Advances in cancer immunotherapy have intensified efforts to identify novel cell surface targets for therapeutic intervention. Surface proteomics provides direct experimental evidence of cell surface localization, rather than relying on *in silico* annotation to retrospectively classify proteins detected by whole-cell proteomics. Therefore, it was intriguing to see that across all cell lines, 56.2% of the surface enriched proteins had an SPC score of 0; suggesting that while they are physically presented on the cell surface, these are not predicted to be there. Our group and others have previously identified RNA-binding proteins (RBPs) like NPM1, snRNP200, and NCL as non-canonical cell surface proteins^3,34–36^, which were well represented in our surfaceome dataset (**Supplementary Figure 8**). In fact, on average 47.7% of the SPC=0 surface proteins were annotated as RBPs (**Figure 4A**).

**Figure 4:**
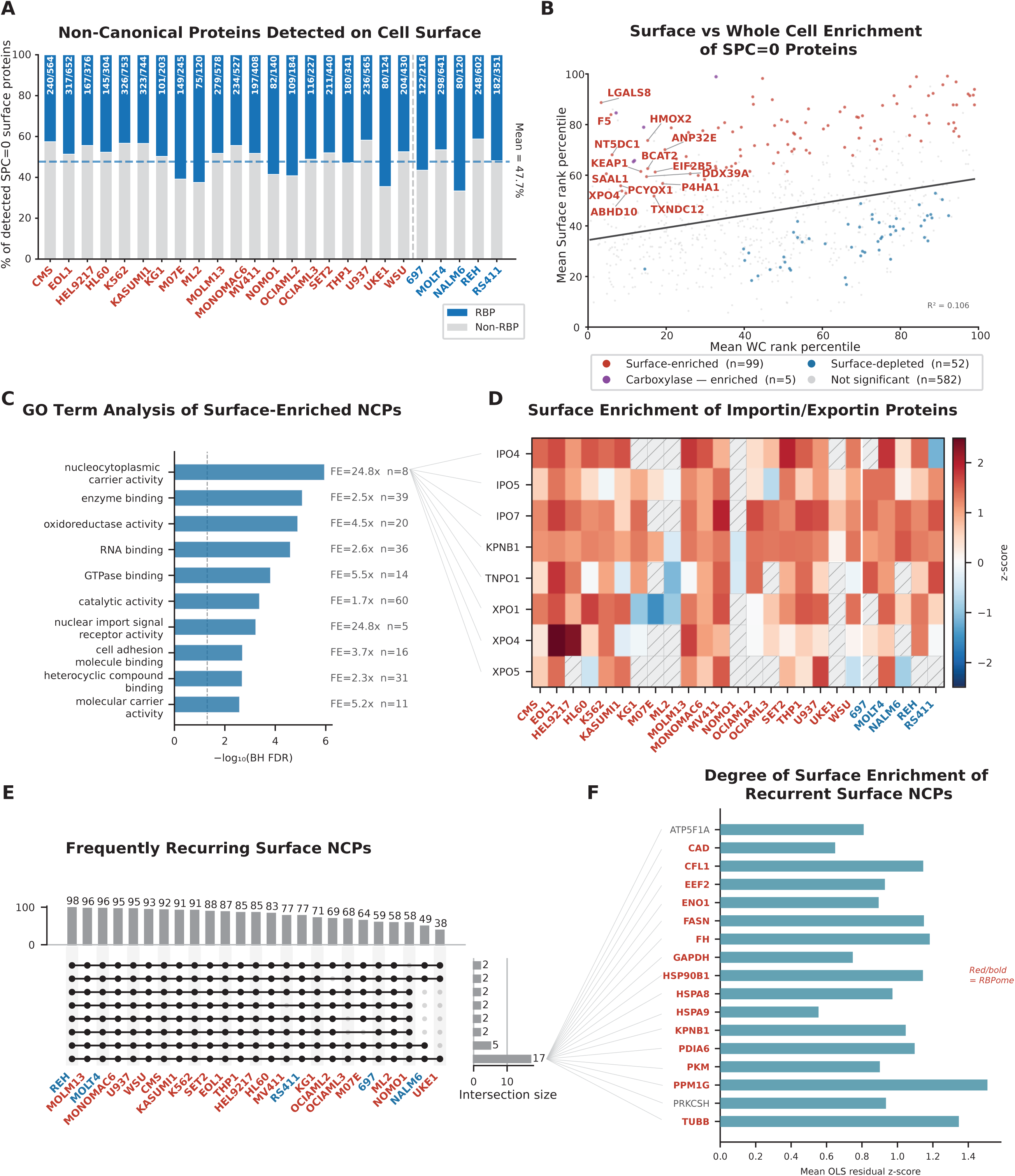
Features of non-canonical cell surface proteins across leukemic cell lines. A. Bar chart showing the fraction of detected SPC=0 surface proteins that are RBPs per cell line, with bar annotations indicating n RBPs / n total SPC=0 detected. The dashed line marks the panel mean of 47.7%. B. Scatter plot of SPC=0 proteins detected in ≥4 cell lines, plotted by its mean whole-cell (x) and mean surface (y) abundance rank percentile across cell lines. Enrichment status was determined by fitting an OLS regression of surface rank percentile on whole-cell rank percentile within each cell line, z-scoring the residuals, and testing the per-protein mean z-score against zero (Wilcoxon signed-rank, BH FDR): surface-enriched (red; FDR<0.05, mean z>0), surface-depleted (blue; FDR<0.05, mean z<0), biotin-dependent carboxylases (purple), or not significant (gray). C. Gene Ontology Molecular Function enrichment analysis of 99 surface-enriched SPC=0 proteins (OLS residual z-score FDR<0.05, mean z>0, excluding biotin-dependent carboxylases) against a background of ∼9,800 proteins detected across whole-cell and surface proteomes. Bars show –log,。(BH-corrected FDR) for the top 10 significant MF terms (Fisher’s exact test); fold-enrichment (FE) and study gene count (n) are annotated to the right of each bar. D. Per-cell-line OLS residual z-scores for nucleocytoplasmic carrier activity genes (GO:0005487) detected in the surface proteome, displayed as a heatmap across 25 cell lines (myeloid, red labels; lymphoid, blue labels; separated by white divider; hatched cells indicate genes not detected in that cell line. E. UpSet plot showing the intersectional detection pattern of surface-enriched SPC=0 proteins (OLS residual z-score FDR<0.05, mean z>0, excluding carboxylases) across all cell lines (myeloid in red, lymphoid in blue). F. The surface-enriched SPC=0 proteins detected in all 25 cell lines, with RBPome members labeled in bold red, OLS residual z-score plotted.

Owing to the paired whole cell proteomics, we could correlate whole cell, and surface proteome ranks of SPC=0 proteins, to identify proteins that are enriched on the cell surface relative to their whole cell abundance. In general, there is a direct relationship between surface and whole cell ranks (**Supplementary Figure 8**). Interestingly, when looking at proteins with disproportionately high representation on the cell surface as compared to the whole cell proteome across all lines, we see many carboxylase proteins enriched (**Figure 4B**). Because several carboxylases are endogenously biotinylated, they likely represent intracellular contaminants captured by streptavidin rather than bona fide cell-surface-enriched proteins, a known phenomenon with biotin capture.

Excluding carboxylases, we identify 99 surface-enriched non-canonical proteins across all cell lines (**Figure 4B**). GO-term analysis revealed RNA binding activity to be enriched among these proteins as expected. Interestingly, nucleocytoplasmic carrier activity was one of the most significant hits (**Figure 4C**). Nucleocytoplasmic carrier activity was represented predominantly by importin and exportin proteins (**Figure 4D**), consistent with prior reports of noncanonical cell-surface localization of nuclear transport proteins.^37,38^ However, this number of related proteins suggests that this class may play a specific role in cell surface biology. Finally, we explored how many of the surface enriched non-canonical proteins were conserved across cell lines. We found that 17 SPC=0 proteins were shared across the surface proteomes of all 25 cell lines (**Figure 4E**), of which 15 are annotated as RNA-binding proteins (**Figure 4F**). This analysis highlights that non-canonical surface proteins, like RBPs, represent a standard feature of the cell surface.

## Discussion

Here we present a comprehensive proteomic atlas of commonly used leukemic cell lines. This dataset serves as a resource to the broader hematologic malignancy community to understand the molecular underpinnings of leukemia and identify potential therapies. Given the extensive transcriptomic characterization of these cell lines, we were also able to make broader observations that highlight the strengths and weakness of these techniques.

RNA sequencing cannot always serve as a reliable surrogate for protein abundance. Although RNA and protein abundance were generally positively correlated, substantial discordance was observed for specific protein classes. Histone proteins were disproportionately abundant by proteomics relative to RNA, whereas mitochondrial genes were prominent among transcripts with comparatively lower protein abundance. These differences may reflect biological factors such as protein stability, turnover, and post-transcriptional regulation, but technical features of both platforms likely also contribute. For example, RNA-sequencing library preparation may influence quantification of histone transcripts, while hydrophobic mitochondrial membrane proteins can be challenging to solubilize, digest, and detect by mass spectrometry. Thus, discordance between RNA and protein abundance should not necessarily be interpreted as evidence of post-transcriptional regulation alone.

Conversely, our data also highlight the greater analytical sensitivity of RNA sequencing, with many low-abundance transcripts detected at the RNA level but not by proteomics, as expected from the amplification inherent to sequencing-based approaches. Owing to the extensive transcriptomic characterization available through the CCLE, the relationships defined here may help inform interpretation of transcriptomic datasets from additional leukemia cell lines, while also identifying contexts in which RNA abundance may not reliably predict protein abundance.

Pairing whole cell proteomics and surface enrichment highlights the complementary utility of these approaches, enabling comprehensive proteome profiling while providing surface specificity. Our work shows two beneficial features of this approach. First, we see that across all cell lines, there are many proteins that are only detected by surface proteomics. This is likely a product of low relative abundance of surface proteins, thus requiring an enrichment strategy to be detected. Second, surface enrichment provides a direct topologic assessment. In general, after whole cell proteomics is performed, in silico predictions and prior literature are used to assign cellular localization to detected proteins. However, work from our group and others have shown that the cell surface contains proteins beyond those with transmembrane domains, GPI-linked anchors, or other dogmatic surface protein features^34^. This resource highlights the broader diversity of the cell surface, including the abundant presence of RNA-binding proteins.

Several limitations should be considered when interpreting the surface-enriched dataset. Cell-surface biotinylation is an enrichment rather than a true fractionation strategy, and proteins detected in this fraction cannot uniformly be assumed to reside on the extracellular surface. Proteomic false-discovery thresholds establish confidence in protein identification but do not distinguish bona fide surface proteins from intracellular proteins captured through nonspecific labeling, cell lysis, or carryover. Accordingly, proteins lacking canonical surface annotations, including SPC=0 and RNA-binding proteins, likely represent a mixture of technical background and proteins with potentially noncanonical surface localization. Orthogonal validation is therefore required before individual candidates are considered therapeutic cell-surface targets. Our use of stringent SPC=0 and SPC=4 categories further prioritize specificity over sensitivity, and proteins with intermediate SPC scores may include additional canonical or noncanonical surface proteins. A second limitation is that all surface-enrichment strategies introduce method-specific biases. Our approach relies on chemical biotinylation of accessible lysine residues, such that lysine abundance and accessibility may influence protein recovery. Consistent with this, predicted surface proteins not detected by surface capture tended to have lower lysine abundance. Alternative approaches, such as periodate oxidation, introduce different biases because capture depends on the presence and composition of sialylated glycans ^39–41^. Thus, no single enrichment strategy is likely to provide a complete representation of the cell-surface proteome, and the most rigorous characterization may require complementary approaches incorporating biotinylation, glycan-based enrichment, and non-affinity-based membrane fractionation.

Beyond providing a proteomic resource, aggregation across cell lines enabled identification of genotype-associated surface proteins. For instance, owing to the large number of *KMT2A*-rearranged cell lines in this dataset, we were able to identify various surface proteins that appear to be upregulated in this population. Of these, CD70 is particularly interesting, as it is currently being developed for CAR-T and ADC therapy. For both modalities, theoretically, increased surface expression should translate to better efficacy and improved therapeutic index. Together, our mass spectrometry findings and corroborating transcriptomic data from primary patient samples nominate CD70 as a therapeutic opportunity in *KMT2Ar* leukemia and support evaluation of CD70-directed therapies in this genetically defined subgroup.

Overall, this study aims to provide a comprehensive dataset for the broader leukemia community and demonstrates the complementary information obtained from whole-cell and surface-enriched proteomics. Integration of these datasets with genomic and transcriptomic annotations enables identification of genotype-associated candidate surface targets, exemplified by CD70 in *KMT2A*-rearranged leukemia, while also revealing a broader set of proteins not captured by conventional surface predictions. Systematic validation of these candidates may expand the repertoire of leukemia-associated antigens available for antibody-, antibody-drug conjugate-, and cellular immunotherapy development.

## Methods

### Cell lines

The leukemia cell-line panel comprised 25 lines, including 20 AML or AML-like myeloid leukemia models and 5 ALL models. The AML panel included fusion-defined subgroups such as KMT2A-rearranged AML, CBFA2T3::GLIS2-rearranged AML, RUNX1::RUNX1T1 AML, FIP1L1::PDGFRA eosinophilic leukemia, and BCR::ABL1-positive myeloid blast-crisis models, as well as mutation-defined models including FLT3-ITD, NPM1-mutant, JAK2 V617F, and RAS-pathway altered lines. The ALL panel included B-ALL and T-ALL models, including *ETV6::RUNX1*, *KMT2A::AFF1*, TCF3::PBX1, and T-ALL cell lines. Cell lines were obtained from the “American Type Culture Collection” (ATCC) or the “Deutsche Sammlung von Mikroorganismen und Zellkulturen” (DSMZ) and cultured in RPMI-1640™ (for HEL 92.1.7, SET-2, CMS, M07e, THP-1, MONO-MAC-6, MV4-11, Kasumi-1, KG-1, NOMO-1, HL-60, UKE-1, WSU, MOLM-13, EOL-1, ML-2, K562, U-937, REH, NALM-6, 697 and MOLT-4) or αMEM™ (for OCI-AML2, OCI-AML3 and RS4;11) + 10% fetal bovine serum (FBS) at 37°C, 5% Co2 atmosphere and 95% humidity. Cell lines were tested and maintained mycoplasma negative throughout all experiments.

### Cell surface protein enrichment

Leukemia cell lines were subjected to cell surface protein enrichment using Sulfo-NHS-SS-biotin labeling. This membrane-impermeable biotinylation reagent selectively labels primary amines on extracellularly exposed proteins, enabling enrichment of cell surface–associated proteins. To minimize labeling of intracellular proteins released from nonviable cells, cell viability of >96% was confirmed prior to labeling. In brief, Leukemia cell lines were subjected to cell surface protein enrichment using Sulfo-NHS-SS-biotin labeling (Pierce Cell Surface Protein Isolation Kit, ID: 89881). For each reaction, 15 million viable cells were washed in PBS and incubated with membrane-impermeable Sulfo-NHS-SS-biotin to selectively label extracellularly exposed primary amines on cell surface proteins. The labeling reaction was quenched, and cells were lysed under mild detergent conditions. Biotinylated surface proteins were captured using NeutrAvidin agarose, washed extensively, and eluted under reducing conditions. Enriched surface proteins were reduced, alkylated and precipitated using methanol/chloroform as previously described ^40^ and the resulting washed precipitated protein was allowed to air dry. Precipitated protein was resuspended in 4 M urea, 50 mM HEPES pH 7.4, followed by dilution to 1 M urea with the addition of 200 mM EPPS, pH 8. Proteins were digested with the addition of LysC (1:50; enzyme:protein) and trypsin (1:50; enzyme:protein) for 12 h at 37 °C. Sample digests were acidified with formic acid to a pH of 2-3 before desalting using C18 solid phase extraction plates (SOLA, Thermo Fisher Scientific). Desalted peptides were dried in a vacuum-centrifuged and reconstituted in 0.1% formic acid for liquid chromatography-mass spectrometry analysis.

Data were collected using an Orbitrap Fusion Lumos mass spectrometer (Thermo Fisher Scientific, San Jose, CA, USA) or an Eclipse mass spectrometer (Thermo Fisher Scientific) coupled with a UltiMate 3000 RSLCnano System. 200ng of peptides were injected and separated on a 50 cm 75 μm inner diameter EasySpray ES903 microcapillary column (Thermo Fisher Scientific), and using a 60 - 70 min gradient of 9 - 30% or 8 - 40% acetonitrile in 1.0% formic acid with a flow rate of 300 - 350 nL/min. Each analysis used a 3 second cycle time data-dependent method. The MS1 data were detected in the Orbitrap over a mass range of m/z 375 – 1325 or 375 - 1500, resolution 120,000, AGC target 4 × 10^5^, 50 - 246 ms maximum injection time, dynamic exclusion of 30 sec, and charge states of 2 - 6. Data-dependent MS2 spectra were isolated in the quadrupole and detected in the Orbitrap with a resolution of 30,000, isolation window of 0.5 m/z, normalized collision energy (NCE) set at 30 - 35%, AGC target 5 x 10^4^ and a 54 ms maximum injection time.

Proteome Discoverer 2.5 (Thermo Fisher Scientific) was used for .RAW file processing and controlling peptide and protein level false discovery rates, assembling proteins from peptides, and protein quantification from peptides. The MS/MS spectra were searched against a Swissprot human database (January 2021) containing both the forward and reverse sequences. Searches were performed using a 10 ppm precursor mass tolerance, 0.03 Da fragment ion mass tolerance, tryptic peptides containing a maximum of two missed cleavages, static alkylation of cysteine (+57.0215 Da), and variable oxidation of methionine (+15.9949 Da), phosphorylation of serine, threonine or tyrosine (+79.9663) and N-terminal acetylation (+42.0106), Met-loss (-131.0405 Da), Met-loss+acetyl (-89.0299 Da). Protein and peptide-spectrum match (PSM) output files were imported into R for downstream analysis using a custom surface proteomics pipeline. PSM-level data were filtered to remove entries with multiple protein accessions or phosphorylation modifications. Protein abundances were scaled by intensity-based absolute quantification (iBAQ) values and normalized at the protein level. Peptide-level data were rolled up to the protein level and filtered to retain proteins identified by at least 2 counts and at least 2 unique peptide sequences across at least 2 replicates.

### Whole cell LFQ quantitative mass spectrometry

Cells were lysed by addition of lysis buffer (8 M Urea, 50 mM NaCl, 50 mM 4-(2-hydroxyethyl)-1-piperazineethanesulfonic acid (EPPS) pH 8.5, Protease and Phosphatase inhibitors) and homogenization by bead beating (BioSpec) for three repeats of 30 seconds at 2400 strokes/min. Bradford assay was used to determine the final protein concentration in the clarified cell lysate. Fifty micrograms of protein for each sample was reduced, alkylated and precipitated using methanol/chloroform as previously described ^40^ and the resulting washed precipitated protein was allowed to air dry. Precipitated protein was resuspended in 4 M urea, 50 mM HEPES pH 7.4, followed by dilution to 1 M urea with the addition of 200 mM EPPS, pH 8. Proteins were digested with the addition of LysC (1:50; enzyme:protein) and trypsin (1:50; enzyme:protein) for 12 h at 37 °C. Sample digests were acidified with formic acid to a pH of 2-3 before desalting using C18 solid phase extraction plates (SOLA, Thermo Fisher Scientific). Desalted peptides were dried in a vacuum-centrifuged and reconstituted in 0.1% formic acid for liquid chromatography-mass spectrometry analysis.

Data were collected using a TimsTOF HT (Bruker Daltonics, Bremen, Germany) coupled to a nanoElute2 LC pump (Bruker Daltonics, Bremen, Germany) via a CaptiveSpray nano-electrospray source. 200 ng of peptides were injected and separated on a reversed-phase C18 column (25 cm x 75 µm ID, 1.6 µM, IonOpticks, Australia) containing an integrated captive spray emitter. Peptides were separated using a 50 min gradient of 2 -30% buffer B (acetonitrile in 0.1% formic acid) with a flow rate of 250 nL/min and column temperature maintained at 50 °C.

The TIMS elution voltages were calibrated linearly with three points (Agilent ESI-L Tuning Mix Ions; 622, 922, 1,222 *m/z*) to determine the reduced ion mobility coefficients (1/K0). To perform diaPASEF, we used py_diAID ^41^, a python package, to assess the precursor distribution in the *m/z*-ion mobility plane to generate a diaPASEF acquisition scheme with variable window isolation widths that are aligned to the precursor density in m/z. Data was acquired using twenty cycles with three mobility window scans each (creating 60 windows) covering the diagonal scan line for doubly and triply charged precursors, with singly charged precursors able to be excluded by their position in the m/z-ion mobility plane. These precursor isolation windows were defined between 350 - 1250 *m/z* and 1/k0 of 0.75 - 1.3 V.s/cm^2^.

The diaPASEF raw files were processed using library-free analysis in DIA-NN 1.8 ^42^, searched against the Swissprot human database (August 2025 for M07e, SET-2, UKE-1, NALM6 and OCI-AML3, and January 2021 for the remaining cell lines). Peptide and protein FDR control, protein assembly, and quantification were performed using default directDIA settings, including tryptic digestion with up to two missed cleavages, carbamidomethylation of cysteine as a fixed modification, and precursor Q-value (FDR) cut-off of 0.01. Precursor quantification strategy was set to Robust LC (high accuracy) with RT-dependent cross run normalization. The DIA-NN output file was imported into R and the precursors mapping to multiple unique protein accessions were removed, and remaining precursor intensities summed to the protein level and iBAQ scaled. Proteins were retained if identified by at least 2 counts and at least 2 unique peptide sequences across at least 2 replicates.

### Surface protein annotations

Replicates for whole cell and enrichment data were assessed for quality control based on peptide and protein abundance, enrichment efficiency, and reproducibility. Replicates failing predefined quality-control thresholds were excluded from downstream cross-sample analyses. A curated surface protein reference table was generated by integrating existing protein annotation resources, including Gene Ontology Cellular Component annotations^45^, UniProt transmembrane and secreted protein annotations^46^, Human Protein Atlas subcellular localization^47^, SURFY^16,48^, and additional surfaceome-focused databases^49^. Identified proteins were annotated for subcellular localization using a curated human proteome reference table derived from the Surface Protein Classification (SPC) framework which assigns proteins to categories including transmembrane and extracellular (surface) and intracellular and other (background).

### Cross-cell-line analysis and candidate prioritization

Normalized protein abundance values were compared across AML and ALL cell lines to identify shared and differentially expressed surface proteins. Differential expression analyses were performed across disease and molecular subgroups, including AML versus ALL, KMT2A-rearranged AML versus other AML lines, and CBFA2T3::GLIS2-rearranged AML versus other AML lines. Candidate surface targets were prioritized based on abundance, detection frequency, differential expression, surface annotation confidence, and potential therapeutic relevance.

### Primary patient data

Surface protein expression patterns were compared with primary patient data obtained from MLL lab. AML and ALL cases were sent for diagnostic work up to MLL Munich Leukemia Laboratory between 2005 and 2022 and analyzed by WTS and WGS. Patients provided written informed consent in accordance with the Declaration of Helsinki.

## Supporting information

Supplementary_Table_S1_WholeCellRaw

Supplementary_Table_S2_SurfaceRaw

Supplementary_Table_S3_WholeCellCompiled

Supplementary_Table_S4_SurfaceCompiled

## Acknowledgments

This work was supported by grants from the Rita Allen Foundation (R.A.F.), the G. Harold and Leila Y. Mathers Charitable Foundation (R.A.F.), Cancer Research Institute (CRI14279; R.A.F.), Pew Charitable Trust, Pew-Stewart Scholars Program (38490; R.A.F.), National Institute Environmental Health Sciences of the National Institutes of Health under award number R01ES037423 (R.A.F.), National Cancer Institute R35 CA283977 (K.S.),ASH Scholar Award (F.W., B.M.G.), Sidney Farber Award, 5K99GM154065 (F.W.), National Cancer Institute K12CA087723 (B.M.G), and DFCI FLAIR Award (B.M.G.).

---

**Supplemental Figure 1:**
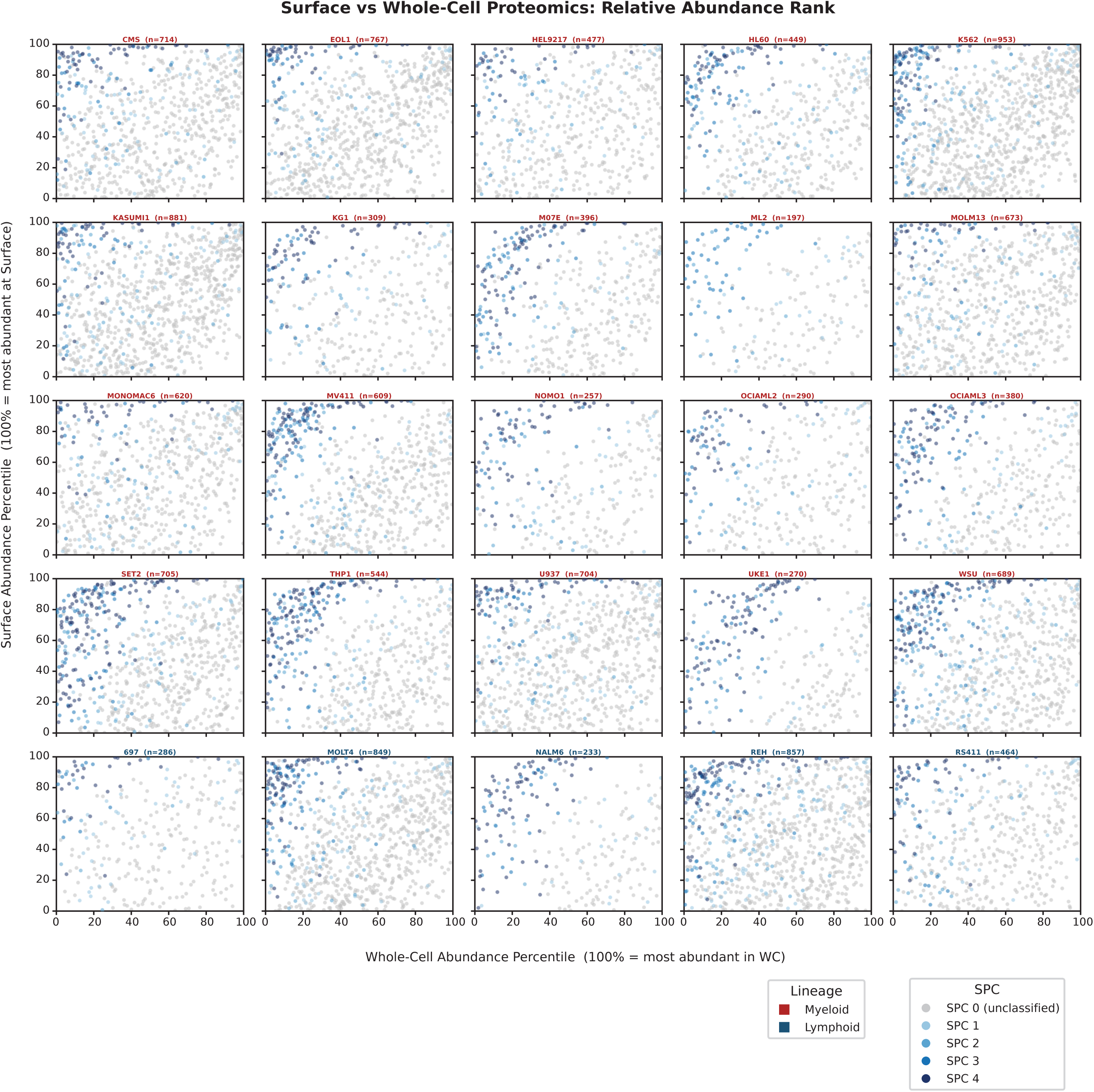
Per cell line correlation of surface and whole cell proteomics. Per-cell-line scatter plots of surface vs whole-cell abundance percentile (100 = most abundant) for proteins detected in both fractions, colored by SPC score (low confidence to SPC 4).

**Supplemental Figure 2:**
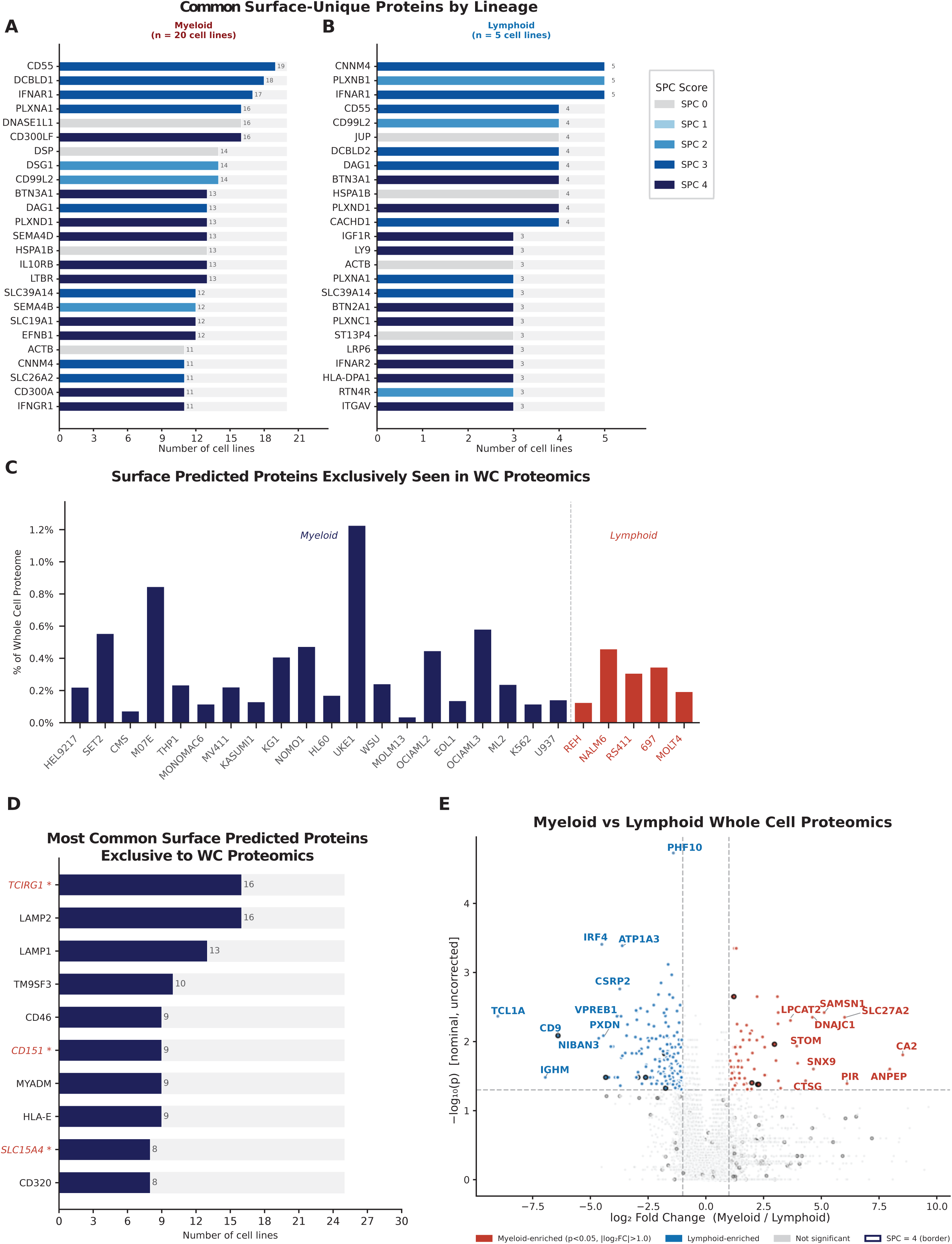
Differential detection of surface annotated proteins in whole cell and surface proteomics. A-B. Top 25 surface-unique proteins (detected in surface proteomics but absent from whole-cell proteomics) ranked by detection frequency across myeloid (A, n=20) and lymphoid (B, n=5) cell lines, colored by SPC score. C. Per-cell-line bar chart showing the percentage of the whole-cell proteome comprised of SPC=4 proteins detected in whole-cell but absent from surface proteomics. D. The 10 most frequently missed SPC=4 proteins in surface proteomics ranked by number of cell lines in which they were missed; asterisks denote proteins never detected in any surface preparation across the panel. E. Volcano plot comparing whole-cell proteomics between myeloid and lymphoid cell lines, using two-sided Mann-Whitney U with a 1 PPM pseudocount for log2 fold change (nominal p < 0.05, |log2FC| > 1.0, uncorrected), with the top proteins by Manhattan distance labeled.

**Supplemental Figure 3:**
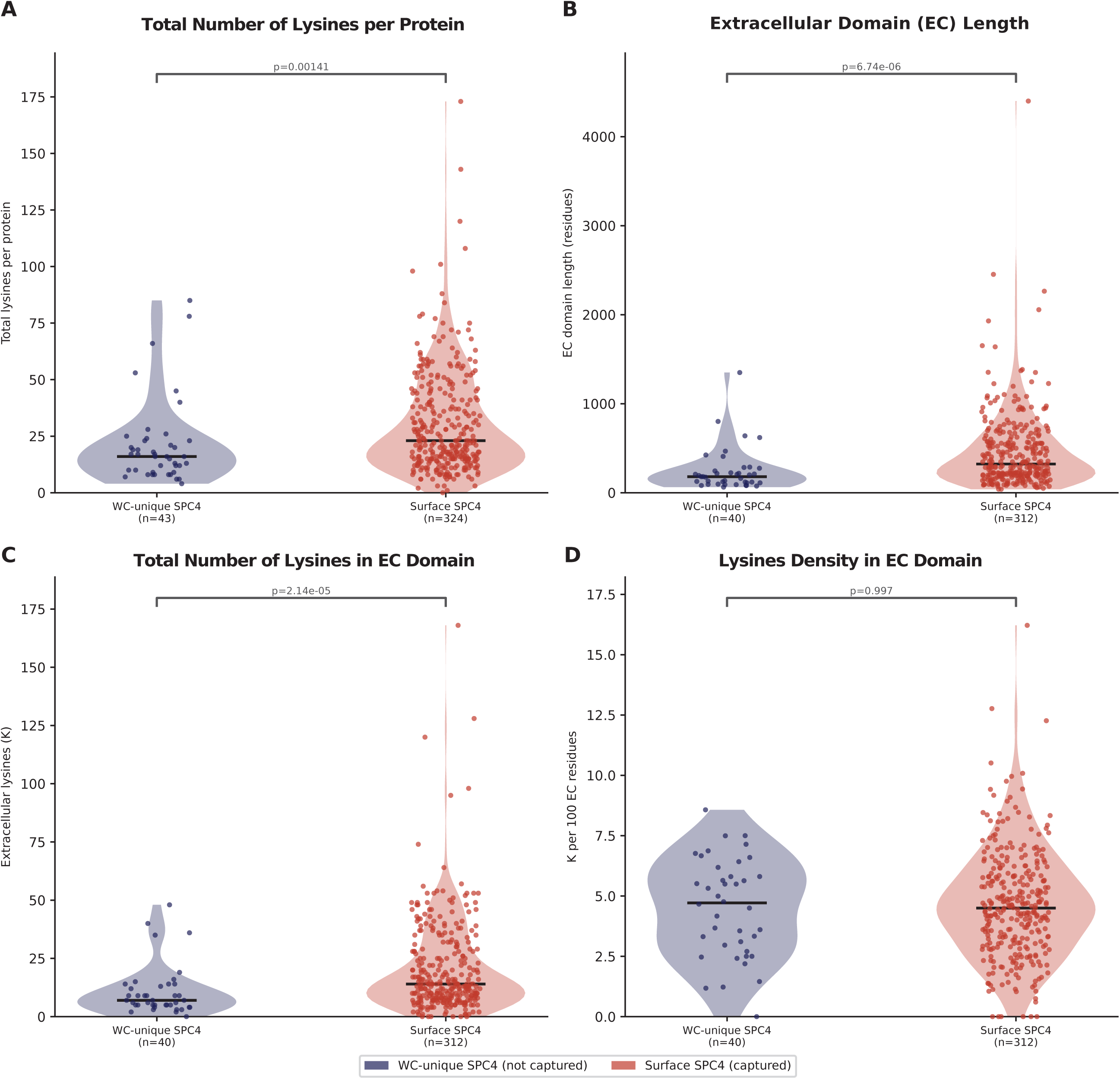
Surface biotinylation is correlated with protein lysine content. A-D. Violin plots comparing lysine content between SPC=4 proteins detected only in whole-cell (WC-unique, not surface captured) and those detected in surface proteomics, using Mann-Whitney U; A shows total lysines per full protein sequence, B shows extracellular (EC) domain length, C shows EC lysine count (biotinylatable residues), and D shows EC lysine density (K per 100 EC residues), with EC annotations derived from UniProt topology features.

**Supplemental Figure 4:**
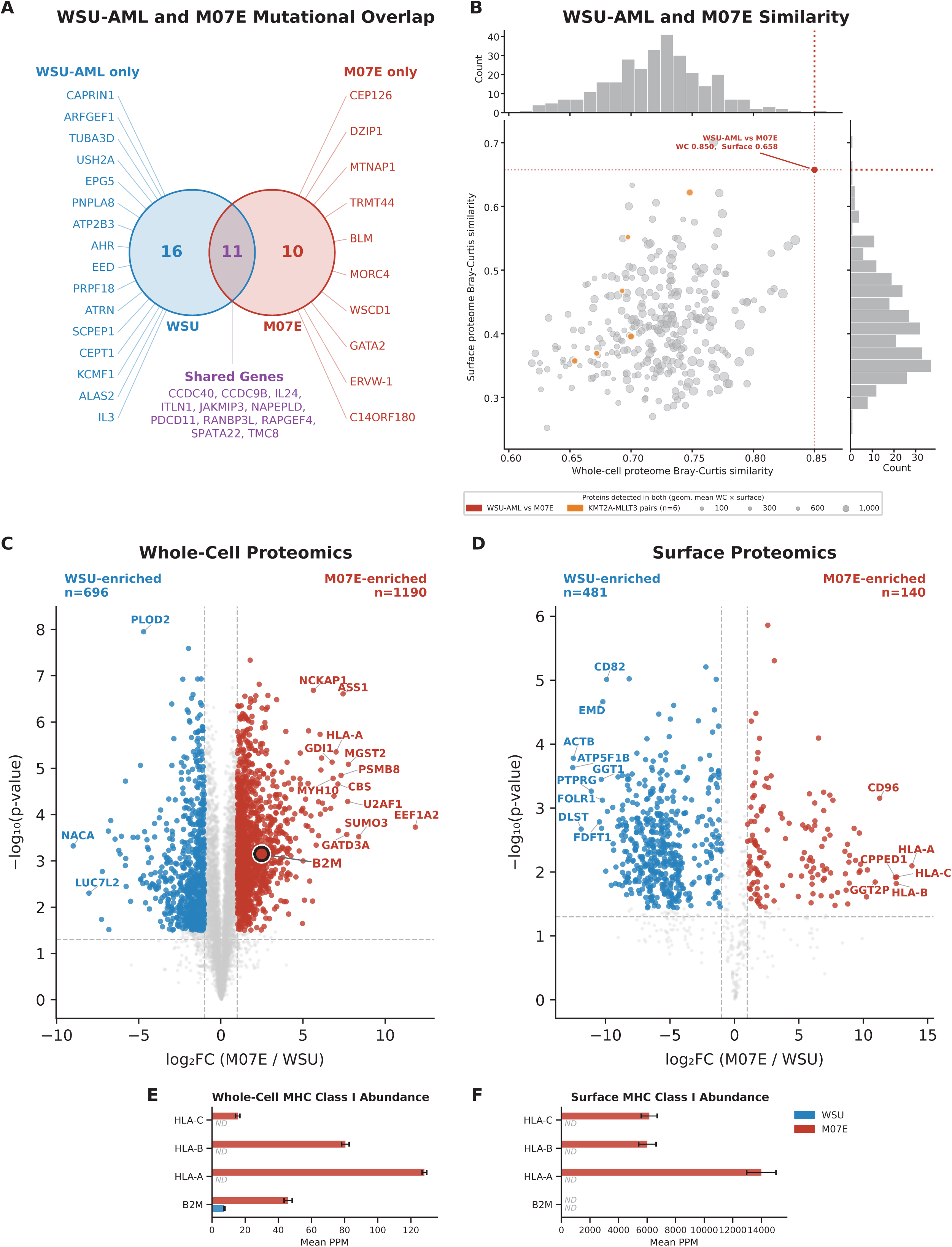
Proteomic comparison of WSU-AML and M07E, both derived from the same patient. A. Venn diagram of HIGH impact somatic mutations identified in WSU-AML and M07E from CCLE. B. Scatter plot of pairwise whole-cell vs surface proteome similarity across all 300 cell line pairs (n=25 cell lines), using Bray-Curtis similarity (presence/absence weighted by relative abundance); bubble size reflects the geometric mean of proteins detected in both proteomes for each pair, and marginal histograms show the distribution of each metric. Cell line pairs with KMT2A-MLLT3 rearrangement are highlighted in orange. C-D. Volcano plots comparing WSU-AML vs M07E in whole-cell (C) and surface (D) proteomics using Welch’s t-test with BH FDR correction (FDR < 0.05, |log2FC| > 1.0), with the top proteins by Manhattan distance per direction labeled. E. Mean PPM of HLA class I heavy chains (HLA-A, HLA-B, HLA-C) and beta-2 microglobulin (B2M) in WSU-AML and M07E whole-cell and surface proteomics. ND = not detected; error bars = SEM across biological replicates.

**Supplemental Figure 5:**
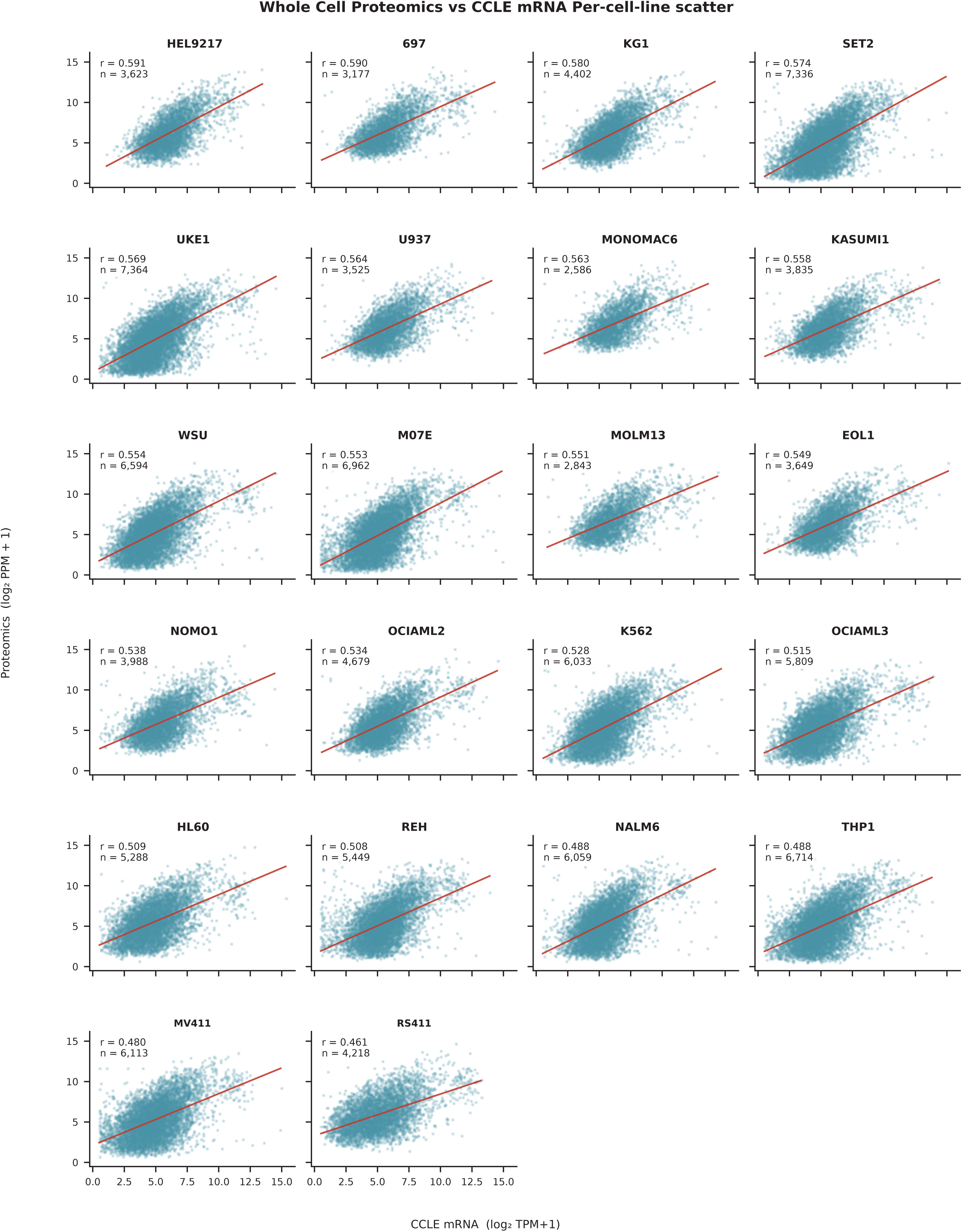
Per-cell-line correlation of RNA and protein abundance. Per-cell-line scatter plots of whole-cell protein abundance (log2 PPM+1) vs CCLE mRNA expression (log2 TPM+1) for genes detected in both, sorted by descending Spearman r; Spearman r and gene count per cell line are shown inset.

**Supplemental Figure 6:**
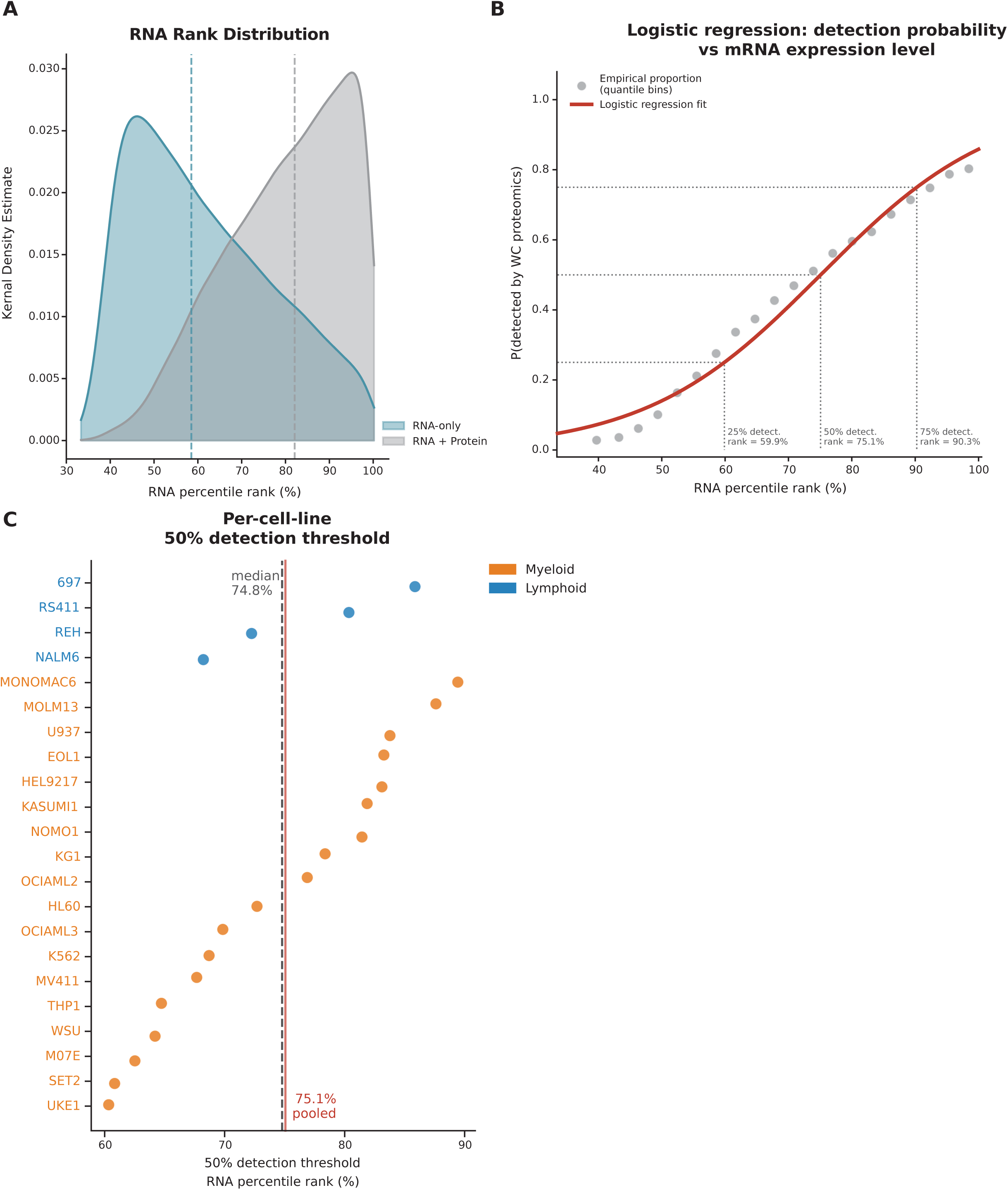
Protein abundance prediction based on RNA expression. A. Kernel density distributions of log2(TPM+1) expression levels for genes detected in both RNA-seq and whole-cell proteomics (RNA + Protein, n=110,223) versus genes detected by RNA-seq alone (RNA-only, n=149,224), pooled across all cell lines; dashed lines indicate group medians. B. Logistic regression modeling the probability of proteomic detection as a function of mRNA expression level, where points represent empirical detection proportions within quantile-binned (20 bins) RNA expression bins (point size proportional to bin size). Dotted lines annotate the log2(TPM+1) thresholds corresponding to 25%, 50%, and 75% probability of proteomic detection. C. Per-cell-line log2(TPM+1) RNA level at which a gene has a 50% probability of proteomic detection, derived from individual logistic regression models; dashed line indicates the median threshold across cell lines.

**Supplemental Figure 7:**
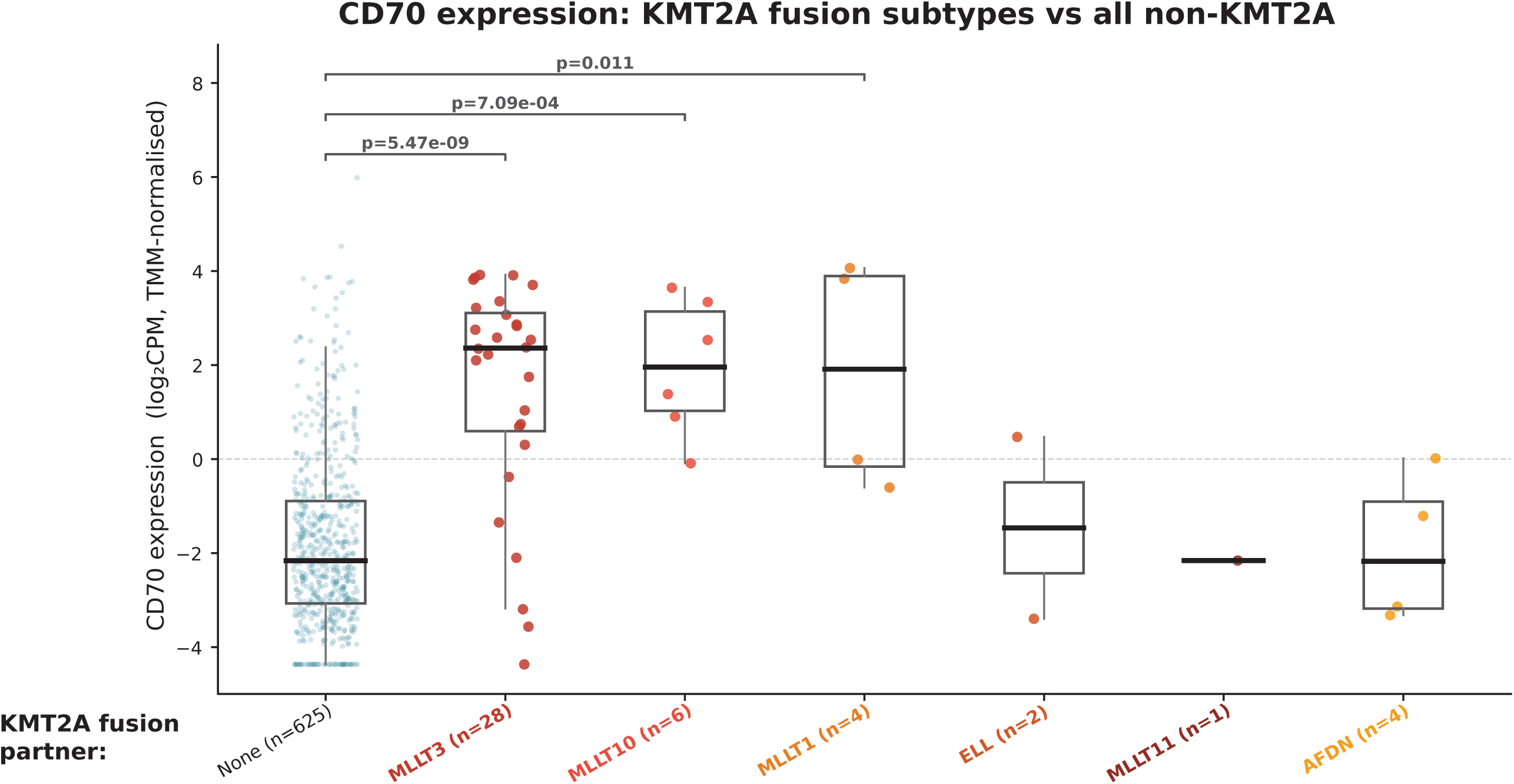
CD70 expression stratified by KMT2A fusion partners. Comparison of CD70 expression obtained from primary patient samples (MLL) across various KMT2A fusions and compared to KMT2A-wildtype samples (blue).

**Supplemental Figure 8:**
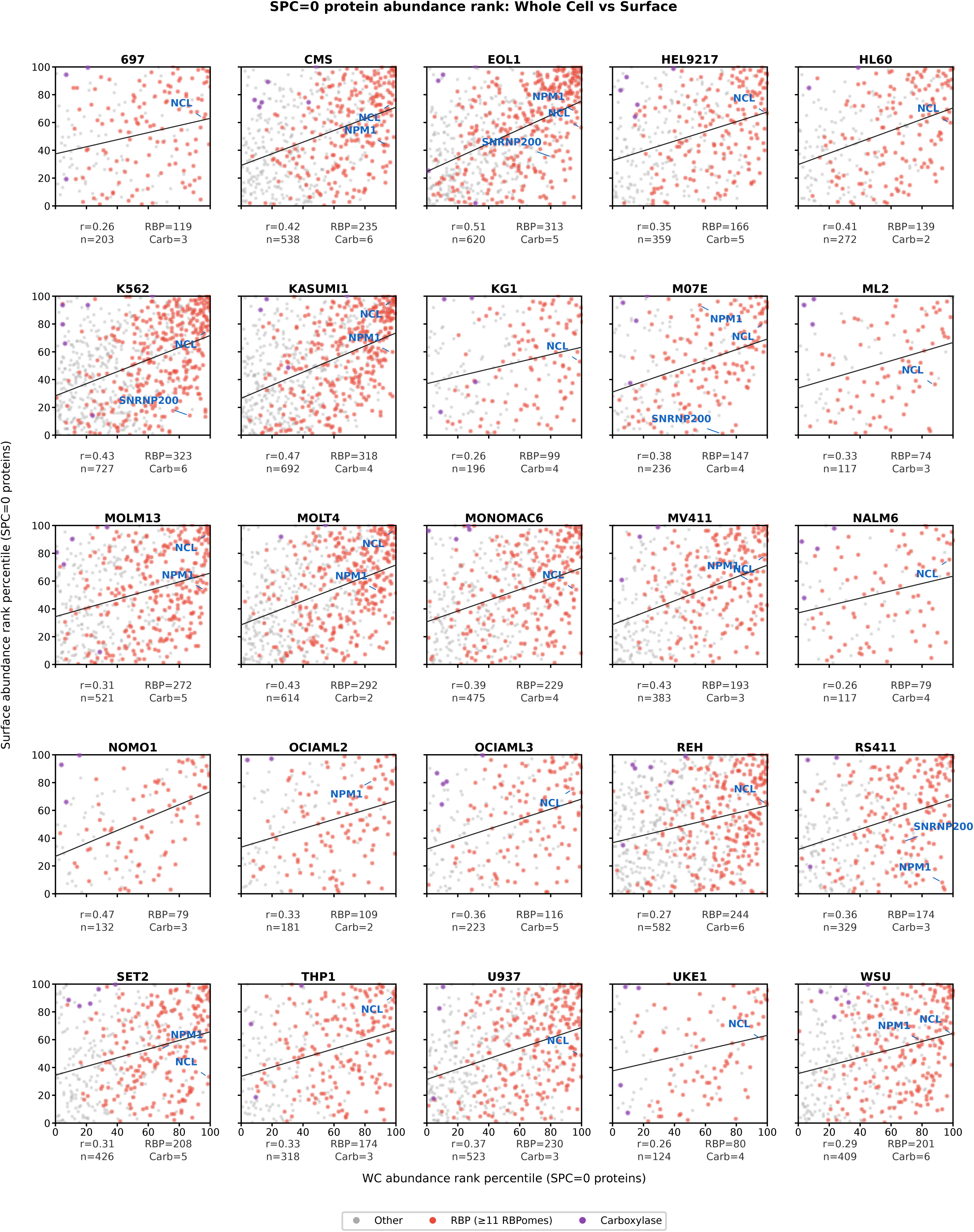
Whole cell and surface proteome relationships of non-canonical surface proteins. Per-cell-line scatter plots of whole-cell (x) versus surface proteomics (y) abundance rank percentile for all SPC=0 proteins detected in each cell line, with OLS trend line and Spearman r shown. RBP and biotin-dependent carboxylase proteins are highlighted in red and purple, respectively; selected genes of interest (NCL, NPM1, SNRNP200) are annotated where detected.

